# Contribution of rare copy number variants to bipolar disorder risk is limited to schizoaffective cases

**DOI:** 10.1101/406215

**Authors:** Alexander W. Charney, Eli A. Stahl, Elaine K. Green, Chia-Yen Chen, Jennifer L. Moran, Kimberly Chambert, Richard A. Belliveau, Liz Forty, Katherine Gordon-Smith, Phil H. Lee, Evelyn J Bromet, Peter F Buckley, Michael A Escamilla, Ayman H. Fanous, Laura J Fochtmann, Douglas S. Lehrer, Dolores Malaspina, Stephen R. Marder, Christopher P. Morley, Humberto Nicolini, Diana O. Perkins, Jeffrey J. Rakofsky, Mark H. Rapaport, Helena Medeiros, Janet L. Sobell, Lena Backlund, Sarah E. Bergen, Anders Juréus, Martin Schalling, Paul Lichtenstein, James A. Knowles, Katherine E. Burdick, Ian Jones, Lisa A Jones, Christina M. Hultman, Roy Perlis, Shaun M. Purcell, Steven A. McCarroll, Carlos N. Pato, Michele T. Pato, Ariana Di Florio, Nick Craddock, Mikael Landén, Jordan W. Smoller, Douglas M. Ruderfer, Pamela Sklar

## Abstract

**Background:** Genetic risk for bipolar disorder (BD) is conferred through many common alleles, while a role for rare copy number variants (CNVs) is less clear. BD subtypes schizoaffective disorder bipolar type (SAB), bipolar I disorder (BD I) and bipolar II disorder (BD II) differ according to the prominence and timing of psychosis, mania and depression. The factors contributing to the combination of symptoms within a given patient are poorly understood.

**Methods:** Rare, large CNVs were analyzed in 6353 BD cases (3833 BD I [2676 with psychosis, 850 without psychosis], 1436 BD II, 579 SAB) and 8656 controls. Measures of CNV burden were integrated with polygenic risk scores (PRS) for schizophrenia (SCZ) to evaluate the relative contributions of rare and common variants to psychosis risk.

**Results:** CNV burden did not differ in BD relative to controls when treated as a single diagnostic entity. Burden in SAB was increased compared to controls (p-value = 0.001), BD I (p-value = 0.0003) and BD II (p-value = 0.0007). Burden and SCZ PRS were higher in SAB compared to BD I with psychosis (CNV p-value = 0.0007, PRS p-value = 0.004) and BD I without psychosis (CNV p-value = 0.0004, PRS p-value = 3.9 × 10^−5^). Within BD I, psychosis was associated with higher SCZ PRS (p-value = 0.005) but not with CNV burden.

**Conclusions:** CNV burden in BD is limited to SAB. Rare and common genetic variants may contribute differently to risk for psychosis and perhaps other classes of psychiatric symptoms.

## INTRODUCTION

Classically conceptualized as an episodic mood disorder with alternating periods of mania and depression, the diagnosis of bipolar disorder (BD) encompasses heterogeneous clinical presentations that vary with respect to symptomatology^1,2^, comorbidity^3^ and longitudinal course^4^. There are 3 diagnoses on the BD spectrum in current classifications of mental illness^5,6^: bipolar I disorder (BD I), bipolar II disorder (BD II) and schizoaffective disorder bipolar type (SAB). The criteria for these diagnoses differ from one another – and from clinically related diagnoses such as schizophrenia (SCZ) and major depressive disorder (MDD) – by nuances in the prominence and timing of manic, depressive and psychotic symptoms. An episode of mania equates to a diagnosis of BD I unless (1) the episode includes psychotic symptoms and (2) there is also a history of psychosis for at least 2 weeks in the absence of mania, in which case SAB is diagnosed. A history of hypomania and depressive episodes equates to a diagnosis of BD II. However, if psychosis occurs during an otherwise hypomanic episode, then the episode is considered manic (and the diagnosis BD I). Psychosis during a depressive episode does not preclude a diagnosis of BD II, so long as the individual has never met the criteria for mania. These nuances are subject to change across versions of the same system of classification^5,7,8^ and the factors determining the combination of symptoms that occur in a given patient remain poorly understood.

BD genetic risk is conferred through many common single-nucleotide polymorphisms (SNPs) of small effect across the genome^9^, many of which also confer risk to clinically related psychiatric conditions^10,11^. The overlap between BD and SCZ is particularly high in this regard, with genetic correlation estimates between the two (r_g_ = 0.6 – 0.7) comparable to estimates between BD I and BD II (r_g_ = 0.7 – 0.8)^9-12^. In contrast, rare variants in particular, rare copy number variants (CNVs) – have not been consistently implicated in risk for BD^13,14,23-26,15-22,^ unlike in SCZ where an increased burden of rare CNVs is well-established^20,22,27,28^. The largest genome-wide study of rare CNVs in BD to date found no differences in burden between approximately 2,600 cases and 8,800 controls^13^. Smaller studies have been inconsistent, with some finding a decreased burden in BD^21,22,29^ and others a modest increase when stratifying cases according to certain clinical criteria. For instance, CNV burden in early-onset BD – a focus of such studies due to the increased CNV burden in neurodevelopmental disorders^25^ has been found by some^15,16,20,26^ but not others^17,21-23^. Specific CNVs implicated in SCZ and neurodevelopmental disorders have been tested for association with BD, and a duplication of 16p11.2 implicated in SCZ^30^ was recently reported to be enriched in BD^13^. Tested as a set rather than individually, these psychiatric CNVs are not significantly enriched in BD^21,22,26^, nor have CNVs in BD consistently been found enriched for particular biological pathways or gene sets^15-17,26^. In total, the evidence that rare CNVs contribute to BD risk broadly is limited.

There is mounting evidence suggesting that the common alleles conferring risk to BD and SCZ act at the symptom level^31,32^, rooting the clinical similarity of BD and SCZ at least partially in common genetic variation. In contrast, the relative absence of rare CNV burden in BD^13^ raises the possibility that this class of variation confers risk to clinical phenomena more commonly associated with SCZ. Such phenomena could include both the nuances in the prominence and timing of psychotic symptoms that formally differentiate SCZ and BD diagnostic criteria^5,6^, as well as non-diagnostic features such as differences in cognitive deficits^33^ and clinical course that historically formed the basis for the dichotomization of BD and SCZ^34,35^. Indeed, studies of psychiatric CNVs in the general population have demonstrated an effect on cognitive performance^33,36^. Profiling rare CNVs and common risk alleles in BD cases stratified by granular clinical data would provide the opportunity to more directly test whether these classes of genetic variation make differential contributions to particular psychiatric traits. To our knowledge, such studies are lacking.

Here, we present results on a genome-wide study of CNVs in BD (6,353 cases and 8,656 controls), approximately a 2.5-fold increase in case sample size from the previous largest such study^13^. In addition to a comprehensive assessment of genome-wide CNV burden between BD and controls, we assess the contribution of rare CNVs and common SCZ risk alleles to risk of psychosis, a clinical phenomenon that differentiates BD subtypes from one another and from SCZ.

## METHODS AND MATERIALS

### Sample Description

The International Cohort Collection for Bipolar Disorder (ICCBD) includes BD cases and unaffected controls from the Sweden Bipolar Disorder Cohort (SWEBIC), the Bipolar Disorder Research Network (BDRN) in the United Kingdom, and the Genomic Psychiatry Consortium (GPC) from the University of Southern California. Full ICCBD sample descriptions have been previously reported in a genome-wide association study (GWAS)^12^. The BDRN controls were collected as part of the Wellcome Trust Case Control Consortium; half were utilized in a genome-wide CNV burden analysis with a set of BD cases not in the current study^22^, and the other half in a separate genome-wide CNV analysis^13^. The subset of the SWEBIC cases and controls genotyped on the Affymetrix platform were in a previous report of genome-wide CNV burden in BD^20^. Genome-wide CNV burden has not been reported before for the GPC cohort or for the SWEBIC cases and controls genotyped on the Illumina platform (45% of ICCBD cases in this study).

### Phenotyping methods

Full descriptions of the approaches utilized in the phenotyping of the ICCBD cohorts have been reported previously^12,37^. For some analyses in this report, clinical variables beyond case-control status were included from all 3 ICCBD sites, including age of onset, history of psychosis and family history. Age of onset was defined as the age at which first symptoms, impairment or diagnosis occurred. Psychosis was defined as the lifetime presence of hallucinations or delusions. Family history was defined as having any family member with any psychiatric diagnosis. For each variable, a set of standardized numerical values were derived, and site investigators harmonized datasets according to these metrics. This was necessary to facilitate analysis across sites that used different phenotyping approaches (***Supplementary Material***).

### Genotyping and ancestry covariates

Sample collection and genotyping procedures for the ICCBD have previously been reported^12^. In brief, for all ICCBD sites DNA was extracted from peripheral blood samples that had been collected and stored at −20°C. Samples were then genotyped at the Broad Institute, and genotypes were called using either Birdsuite (Affymetrix) or BeadStudio (Illumina). Genotypes were generated as sufficient numbers of samples accumulated from field work. Ancestry covariates were derived from the genotyping data through multidimensional scaling (MDS) analysis on genome-wide identity-by-descent distances calculated for all pairs of individuals. Full details on the quality control procedures implemented to derive the genotype calls utilized in this report have been previously described^12^.

### CNV calling and quality control

Rare CNVs were identified using the Birdseye program in Birdsuite^38^, which is based on a hidden Markov model. For each CNV, a logarithm of the odds ratio (LOD) score was generated that describes the likelihood of a CNV relative to no CNV over a given interval including flanking sequences. Only subjects who passed quality control filters in an earlier GWAS of the same individuals^12^ were considered for CNV analyses. CNVs were excluded if any of the following criteria were met: LOD score < 10, number of probes < 10, probe density of < 1 per 20 kilobases (KB), frequency in ICCBD > 1%, or location within a region known to contain common CNVs or large genomic gaps (e.g., centromeres). If in a given individual the distance between two CNVs was less than 20% of their combined size, they were considered artificially split by the calling algorithm and combined into a single event. For the BDRN cohort, only genomic regions covered in both cases and controls were retained in order to reduce batch effects resulting from cases and controls being genotyped on different Illumina arrays (***Supplementary Material***). Subjects were removed for having total CNV number greater than two standard deviations different from the mean number of CNVs in the cohort (prior to applying filters for CNV frequency). These quality control checks were performed separately for the SWEBIC Affymetrix, SWEBIC Illumina, BDRN, and GPC cohorts (***Table 1***).

**Table 1:** Sample characteristics for ICCBD CNV analyses.

| Cohort | Ethnicity | Chip | Batch<sup>a</sup> | Cases | Controls | BD I |  |  | BD II | SAB | NOS |
| --- | --- | --- | --- | --- | --- | --- | --- | --- | --- | --- | --- |
|  |  |  |  |  |  | All | Psychosis | No psychosis |  |  |  |
| SWEBIC | European | Affymetrix 6.0 | Wave3 | 0 | 499 | 0 | 0 | 0 | 0 | 0 | 0 |
|  |  | Affymetrix 6.0 | Wave4 | 917 | 1170 | 538 | 355 | 162 | 114 | 20 | 245 |
|  |  | Omni Express | Wave6 | 1,356 | 1140 | 597 | 355 | 199 | 500 | 30 | 229 |
| BDRN | European | Combo Chip | - | 1,084 | 0 | 677 | 389 | 185 | 361 | 34 | 12 |
|  |  | Omni Express | - | 1,493 | 0 | 991 | 694 | 161 | 426 | 57 | 19 |
|  |  | Illumina 1.2M | 58BC | 0 | 2428 | 0 | 0 | 0 | 0 | 0 | 0 |
|  |  | Illumina 1.2M | NBS | 0 | 2149 | 0 | 0 | 0 | 0 | 0 | 0 |
| GPC | European-American | Omni Express | - | 1,503 | 1,270 | 1,030 | 883 | 143 | 35 | 438 | 0 |
| **ICCBD** |  |  |  | **6,353** | **8,656** | **3,833** | **2,676** | **850** | **1,436** | **579** | **505** |

<sup>a</sup>CNV analyses including Swedish samples from waves 2-4 have been previously reported in Bergen et al. CNV results for SWEBIC controls from all waves have been reported in Marshall et al. CNV analyses for BD using subsets of the BDRN controls have been reported in Grozeva et al 2010 and Green et al 2016. BDRN cases were included in analyses by Green et al 2016.

Unless otherwise specified, burden analyses were restricted to autosomal CNVs >100kb. Two events were considered equivalent for the purposes of defining frequency if one overlapped the other by at least 50%. In the context of burden analyses, we use the term “CNV” to refer to the combined set of deletions and duplications, and “singleton CNVs” were defined as any event that occurred once in the full ICCBD case-control cohort without consideration of whether the event was a deletion or a duplication. Specific singleton deletions and duplications were defined after first filtering the dataset for that type of event. As such, not all singleton deletions and duplications are in the singleton CNV group.

### CNV burden tests

For our primary analyses, we defined CNV burden in 3 ways: the number of CNVs occurring per individual (the CNV number); the number of genes lying within CNVs per individual (the CNV gene count); the total distance covered by CNVs. We elected to focus on these 3 classes of burden because there is no clear class of burden most relevant to BD and these classes significantly differed between cases and controls in the largest SCZ CNV study to date^27^. We stratified CNVs by 3 types: deletions only, duplications only, deletions and duplications (or “CNVs”); by 2 sizes: over 100KB and over 500KB; and by 2 frequencies: singletons and those occurring in less than 1% in the ICCBD (a frequency of 6.7 × 10^−5^). This led to 36 tests between each pair of phenotypes we compared, of which there were 7: (1) BD cases to controls, (2) BD I cases to controls, (3) BD II cases to controls, (4) SAB cases to controls, (5) BD I cases to BD II cases, (6) BD I cases to SAB cases, and (7) BD II cases to SAB cases. Therefore, in total, there were 252 tests in our primary assessment of CNV burden.

Previous studies of CNV burden in BD have reported nominally significant results (p-value < 0.05) for tests where the definition of burden fell outside the scope of the 252 tests in our primary burden assessment. Through manual curation of the literature, we identified 34 unique associations (2 were observed in 2 separate studies). We were able to test 27 of these in the ICCBD data (for the other 7, the original study included either SCZ cases or BD parent-child trios), of which 21 included a burden class not assessed in our primary 252 tests. In some instances, the dataset used in the original paper overlapped that used in this report, in which case the overlapping samples were excluded from the test.

We also tested ICCBD CNVs, filtered for size over 100KB and frequency <1%, for enrichment of sets of CNVs previously identified in studies of BD, SCZ or neurodevelopmental disorders. The BD CNV set (16 deletions, 14 duplications) was derived by merging overlapping autosomal *de novo* CNVs from 3 previous studies of BD trios^16,17,24^. The SCZ CNV set was comprised of autosomal CNVs with suggestive evidence for association in a meta-analysis of over 20,000 SCZ cases and 20,000 controls (11 deletions, 8 duplications)^27^. The neurodevelopmental CNV set was derived from a list curated for a previous report^17^ for which CNVs overlapping those in the SCZ set had been removed (27 deletions, 18 duplications). In order for a CNV in the test set to be considered overlapping with an ICCBD CNV, the ICCBD CNV was required to cover at least 50% of the test CNV and be of the same CNV type (i.e., deletion or duplication).

All tests were performed using permutation in PLINK^39^ controlling for genotyping platform and ICCBD site. Significance was evaluated using 10,000 permutations. The 252 tests in the primary assessment were 2-sided with the exception of 6 tests that had previously been reported as significant. A one-sided test in the direction of the association reported in the original paper was used for these 6 tests as well as for the additional 21 tests following up previous associations and the 3 tests of CNV sets. To account for multiple testing, we considered as study-wide significant any result surpassing correction for 276 tests (252 primary burden tests, 21 tests of previous burden associations, 3 tests of CNV sets). At a 5% false discovery rate (FDR), an empirical p-value below 0.002 was considered study-wide significant.

### Contribution of CNV burden and SCZ PRS to psychosis

Following results from our primary burden analyses, we analyzed CNV burden and loading of common SCZ risk alleles in BD I and SAB cases. BD II was excluded from these analyses to remove effects resulting from known differences in polygenic loading of SCZ alleles across BD subtypes^12^. For these analyses, burden was defined as the number of CNVs greater than 500KB and present in less than 1% of the study sample. We focused on this one burden definition here because it was the only class in our primary assessment of burden where an increase was seen in SAB compared to controls, BD I and BD II (see ***Results***). For these analyses, burden was tested using logistic regression, which returned similar results to permutation but allowed us to include in the model continuously-distributed ancestry covariates and facilitated the calculation of odds-ratios (ORs) for CNV burden^27^. In the regression model, we used phenotype status as the dependent variable and CNV burden as an independent predictor variable. The OR is the exponential of the logistic regression coefficient, and OR > 1 represents increased risk for the “affected” phenotype in the model, which was designated to be the phenotype more clinically similar to SCZ. Using a similar regression model, we carried out polygenic scoring analyses^40^. Quantitative polygenic risk scores (PRS) were computed for each case subject based on the set of SNPs with p-values less than 0.5 in the second SCZ GWAS from the Psychiatric Genomics Consortium (PGC)^41^. PRS analyses excluded ICCBD samples present in the PGC studies and ICCBD SWEBIC Affymetrix cases due to lack of a control cohort once PGC overlaps were removed. Effect sizes for both CNV burden and SCZ PRS were calculated as a t-statistic that is the ratio of the coefficient of the burden or PRS variable and its standard error from a generalized linear regression model equation. As studies of SCZ have consistently demonstrated higher CNV burden in cases compared to controls^27,28^, cases were stratified by clinical dimensions related to SCZ (i.e., psychosis) and 1-sided statistical tests were used evaluating for higher rates in groups with the more SCZ-like phenotype.

## RESULTS

### CNV burden in BD

We assessed genome-wide differences in rare CNV burden between 6,353 BD cases and 8,656 controls (***Table 1)***. After initial filters for size (> 100KB) and frequency (occurring in < 1% of ICCBD), we observed 10,515 CNVs (3,970 deletions and 6,545 duplications). No difference in the CNV number was found between cases and controls (case rate = 0.698, control rate = 0.702, p-value = 0.86). This was true both for deletions (case rate = 0.266, control rate = 0.264, p-value = 0.78) and duplications (case rate = 0.433, control rate = 0.439, p-value = 0.72). Similarly, no differences were observed between cases and controls with respect to the number of genes hit or the total distance covered by CNVs (***Table 2***).

**Table 2:**
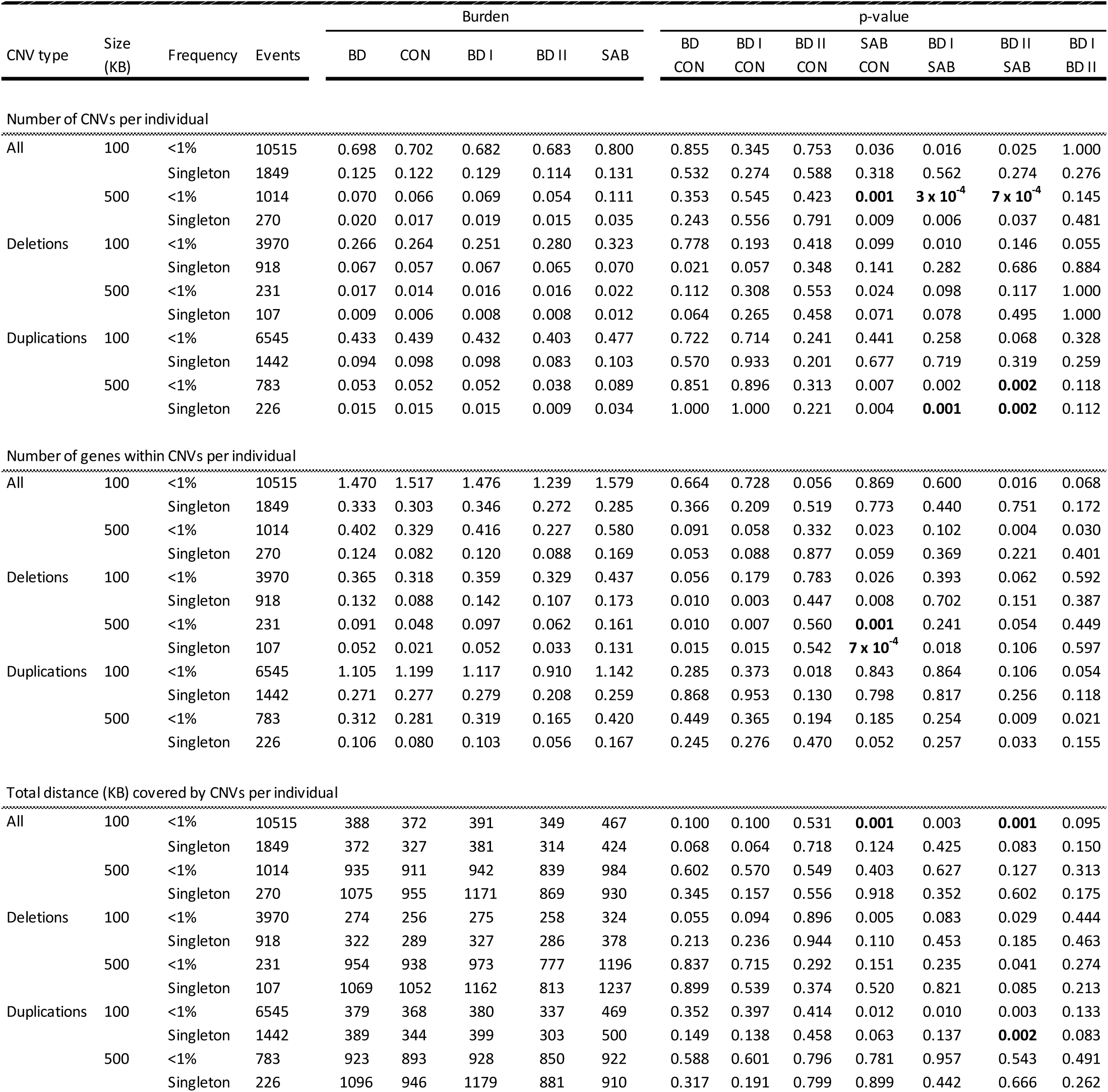
CNV burden in bipolar disorder. Burden metrics are presented for the following groups (N): BD treated as a single diagnostic entity (6353), controls (8656), BD I (3833), BD II (1436) and SAB (579). Three classes of CNV burden were assessed, indicated by the description above the dotted lines. Burden was compared between cases and controls, as well as between case subtypes. P-values are two-sided, uncorrected for multiple testing, and based on 10,000 permutations testing for relative burden between the two groups. CNV type, size, and frequency refer to the filters applied for the test being reported. Events refers to the number of CNVs total observed in the groups being compared for the specified parameters. P-values surpassing study-wide significance are shown in bold. Singletons are those CNVs that occur once within the full ICCBD case cohort when filtered for those greater than 100KB in size. KB=kilobases

Following previous literature showing that rarer and larger CNVs carry increased burden for neuropsychiatric illness^28^, we further filtered CNVs by size (> 500KB) and frequency (those that occur once in the 15,009 ICCBD individuals, a frequency of 6.6 × 10^−5^). No test was significant below our study-wide p-value threshold (***Table*** 2) that accounts for 276 genome-wide burden tests at FDR 5% (see ***Methods***). A report on CNV burden for the SWEBIC Affymetrix sample (917 cases, 1,1670 controls) previously noted a significant increase in the number of CNVs in BD cases compared to controls^20^. To assess whether nominally significant associations (***Table 2***) were driven by these previously reported observations, we repeated these analyses after excluding the SWEBIC Affymetrix cohort and no tests remained nominally significant (***Supplementary Material***).

Previous reports of CNV burden in BD have found nominally significant associations across several classes of burden beyond those assessed above. We curated the literature to identify all previous associations of BD and CNV burden of at least nominal significance (p-value < 0.05) in the initial report. We found 34 unique associations, of which we could test 21 that were not included as part of our primary assessment of burden (see ***Methods***). None of these tests surpassed study-wide significance p-value threshold of 0.002 (***Figure 1; Table 3***).

**Table 3:** Follow-up of previous reports of increased copy number variant burden in BD. Listed are all findings identified from manual curation of literature on rare copy number variant (CNV) burden in BD with a p-value below 0.05 in the original report. Details of the test performed in the original report that was reproduced in ICCBD are described in the test parameter fields and include the phenotypes compared, the definition of burden, and the filters applied for CNV frequency, type and size. Ratios were calculated as the burden in the first phenotype in the comparison field relative to the second phenotype. When the original report did not specify the CNV size studied all CNVs greater than 100KB were included in the ICCBD. Original reports where the test included either SCZ cases or BD trios could not be followed-up in the ICCBD, but are included in this table so as to consolidate all of the previously significant findings in rare CNV studies of BD. BD cases and controls in Bergen (2011) are part of the ICCBD, so for these follow-up tests only the ICCBD samples not in the original report were utilized. The controls in Grozeva (2010) comprise half of the BDRN controls in ICCBD; since there was no case overlap between these studies, all of these controls were included in the follow-up test in the ICCBD. The cases in Green (2015) are the BDRN cases in ICCBD, though these findings were not followed up in ICCBD as they involved SCZ cases. Early-onset was defined as less than 21 in Priebe (2012) and less than 18 in Zhang (2009) and ICCBD. Family history in Malhotra (2011) was defined as having a relative with bipolar disorder (I, II, SAB), schizophrenia, autism, MDD or intellectual disability; in ICCBD, it was defined as having a family member with any psychiatric history. Reported p-values for ICCBD are 1-sided from using 10,000 permutations to test for enrichment in the direction observed in the original report. Asterisks denote nominal significance observed in ICCBD. BD – bipolar disorder; CON – control; EO – early-onset bipolar disorder; FAM – bipolar disorder with a family history of psychiatric illness; SPOR – sporadic bipolar disorder (i.e., no family history of psychiatric illness); SCZ – schizophrenia; SING – singleton; CNV – copy number variant; SCZ – schizophrenia; NR - not reported; NA - not applicable; N - number of individuals for the groups listed in the comparison field

| Test parameters |  |  |  |  | Original report |  |  |  | ICCBD |  |  |
| --- | --- | --- | --- | --- | --- | --- | --- | --- | --- | --- | --- |
| Comparison | Freq | Type | Burden definition | Size (KB) | Author (year) | N | Ratio | p-value | N | Ratio | p-value |
| BD/CON | < 1% | CNV | Gene set enrichment: "psychological disorders" | >=100 | Zhang (2009) | 998/1001 | NR | $6 \times 10^{-6}$ | 6353/8656 | 0.90 | 0.806 |
| BD/CON | < 1% | CNV | Gene set enrichment: "behavior (learning)" | >=100 | Zhang (2009) | 998/1001 | NR | $6 \times 10^{-3}$ | 6353/8656 | 1.04 | 0.603 |
| BD/CON | < 1% | CNV | Number of CNV per individual | >=100 | McQuillin (2011)<sup>a</sup> | 500/500 | 0.82 | 0.0360 | 6353/8656 | 1.01 | 0.700 |
| BD/CON | < 1% | CNV | Number of CNV per individual | >=1000 | Grozeva (2013) | 1650/10259 | 0.58 | 0.0100 | 6353/8656 | 0.84 | 0.962 |
| BD/CON | < 1% | CNV | Number of *de novo* CNV per individual | >=10 | Malhotra (2011) | 185/426 | 4.78 | 0.0090 | NA | NA | NA |
| BD/CON | < 1% | CNV | Number of *de novo* CNV per individual | >=10 | Georgieva (2014) | 768/3208 | 2.15 | 0.0007 | NA | NA | NA |
| BD/CON | < 1% | DEL | Number genes hit by CNV per individual | >=500 | Bergen (2012)<sup>b</sup> | 834/2087 | 2.12 | 0.0390 | 5436/6987 | 1.73 | 0.064 |
| BD/CON | < 1% | DEL | Number of CNV per individual | >=100 | Grozeva (2010) | 1697/2806 | 0.83 | 0.0100 | 6353/8656 | 1.01 | 0.355 |
| BD/CON | < 1% | DEL | Number of CNV per individual | >=1000 | Grozeva (2013) | 1650/10259 | 0.44 | 0.0300 | 6353/8656 | 1.45 | 0.102 |
| BD/CON | < 1% | DEL | Number of CNV per individual | 100-200 | Grozeva (2010) | 1697/2806 | 0.83 | 0.0300 | 6353/8656 | 0.98 | 0.636 |
| BD/CON | < 1% | DEL | Number of CNV per individual | 200-500 | McQuillin (2011)<sup>a</sup> | 500/500 | 0.64 | 0.0390 | 6353/8656 | 1.04 | 0.312 |
| BD/CON | SING | CNV | Number genes hit by CNV per individual | >=100 | Bergen (2012)<sup>b</sup> | 834/2087 | 1.37 | 0.0320 | 5436/6987 | 1.00 | 0.428 |
| BD/CON | SING | DEL | Average size of genic CNVs | NR | Priebe (2012) | 882/872 | 1.90 | 0.0140 | 6353/8656 | 1.04 | 0.346 |
| BD/CON | SING | DEL | Proportion of individuals with a CNV | >=100 | Zhang (2009) | 998/1001 | 1.33 | 0.0070 | 6353/8656 | 1.17 | 0.012 |
| BD/CON | SING | DEL | Number genes hit by CNV per individual | >=100 | Bergen (2012)<sup>b</sup> | 834/2087 | 1.93 | 0.0100 | 5436/6987 | 1.24 | 0.153 |
| BD/CON | SING | DEL | Number of CNV per individual | >=100 | Zhang (2009) | 998/1001 | 1.38 | 0.0100 | 6353/8656 | 1.17 | 0.014 |
| BD/CON | SING | DEL | Number of CNV per individual | >=100 | Bergen (2012)<sup>b</sup> | 834/2087 | 1.35 | 0.0130 | 5436/6987 | 1.12 | 0.150 |
| BD/CON | SING | DEL | Total size of genic CNVs | NR | Priebe (2012) | 882/872 | 1.84 | 0.0140 | 6353/8656 | 1.07 | 0.240 |
| BD/SCZ | < 1% | CNV | Number of *de novo* CNV per individual | >=100 | Georgieva (2014) | 768/1115 | 0.51 | 0.0150 | NA | NA | NA |
| BD/SCZ | < 1% | DEL | Number of exonic CNV per individual | >=1000 | Green (2015) | 2591/6882 | 0.40 | 0.0009 | NA | NA | NA |
| BD/SCZ | < 1% | DUP | Number of exonic CNV per individual | 500-1000 | Green (2015) | 2591/6882 | 0.80 | 0.0450 | NA | NA | NA |
| EO/CON | < 1% | CNV | Proportion of individuals with a genic CNV | NR | Priebe (2012) | 291/872 | 1.26 | 0.0009 | 2235/8656 | 0.96 | 0.529 |
| EO/CON | < 1% | CNV | Number of *de novo* CNV per individual | >=10 | Malhotra (2011) | 107/426 | 6.22 | 0.0060 | NA | NA | NA |
| EO/CON | < 1% | DUP | Proportion of individuals with a genic CNV | NR | Priebe (2012) | 291/872 | 1.34 | 0.0004 | 2235/8656 | 0.97 | 0.275 |
| EO/CON | < 1% | DUP | Number of genic CNV per individual | NR | Priebe (2012) | 291/872 | NR | 0.0240 | 2235/8656 | 1.00 | 0.108 |
| EO/CON | SING | CNV | Proportion of individuals with a CNV | >=100 | Zhang (2009) | NR/1001 | 1.17 | 0.0390 | 2235/8656 | 1.05 | 0.361 |
| EO/CON | SING | CNV | Number of genic CNV per individual | NR | Priebe (2012) | 291/872 | NR | 0.0450 | 2235/8656 | 1.14 | 0.251 |
| EO/CON | SING | DEL | Average size of genic CNVs | NR | Priebe (2012) | 291/872 | 2.65 | 0.0056 | 2235/8656 | 1.20 | 0.148 |
| EO/CON | SING | DEL | Proportion of individuals with a CNV | >=100 | Zhang (2009) | NR/1001 | 1.58 | 0.0010 | 2235/8656 | 1.22 | 0.145 |
| EO/CON | SING | DEL | Number of CNV per individual | >=100 | Zhang (2009) | NR/1001 | 1.54 | 0.0040 | 2235/8656 | 1.25 | 0.106 |
| EO/CON | SING | DEL | Total size of genic CNVs | NR | Priebe (2012) | 291/872 | 2.60 | 0.0084 | 2235/8656 | 1.19 | 0.141 |
| EO/CON | SING | DUP | Proportion of individuals with a genic CNV | NR | Priebe (2012) | 291/872 | 1.45 | 0.0170 | 2235/8656 | 0.98 | 0.613 |
| EO/CON | SING | DUP | Number of genic CNV per individual | NR | Priebe (2012) | 291/872 | NR | 0.0100 | 2235/8656 | 0.97 | 0.675 |
| FAM/CON | < 1% | DUP | Number of genic CNV per individual | >=500 | Malhotra (2011) | 107/426 | 1.83 | 0.0300 | 3072/8656 | 1.09 | 0.066 |
| SPOR/CON | < 1% | CNV | Number of *de novo* CNV per individual | >=10 | Malhotra (2011) | 78/426 | 4.22 | 0.0190 | NA | NA | NA |
| SPOR/FAM | < 1% | CNV | Number of *de novo* CNV per individual | >=10 | Georgieva (2014) | 318/50 | 0.31 | 0.0390 | NA | NA | NA |

<sup>a</sup>Sample sizes approximated from earlier studies of the same cohort
<sup>b</sup>This study included the ICCBD SWEBC Affymetrix cohort, thus our replication test excluded this cohort

**Figure 1.**
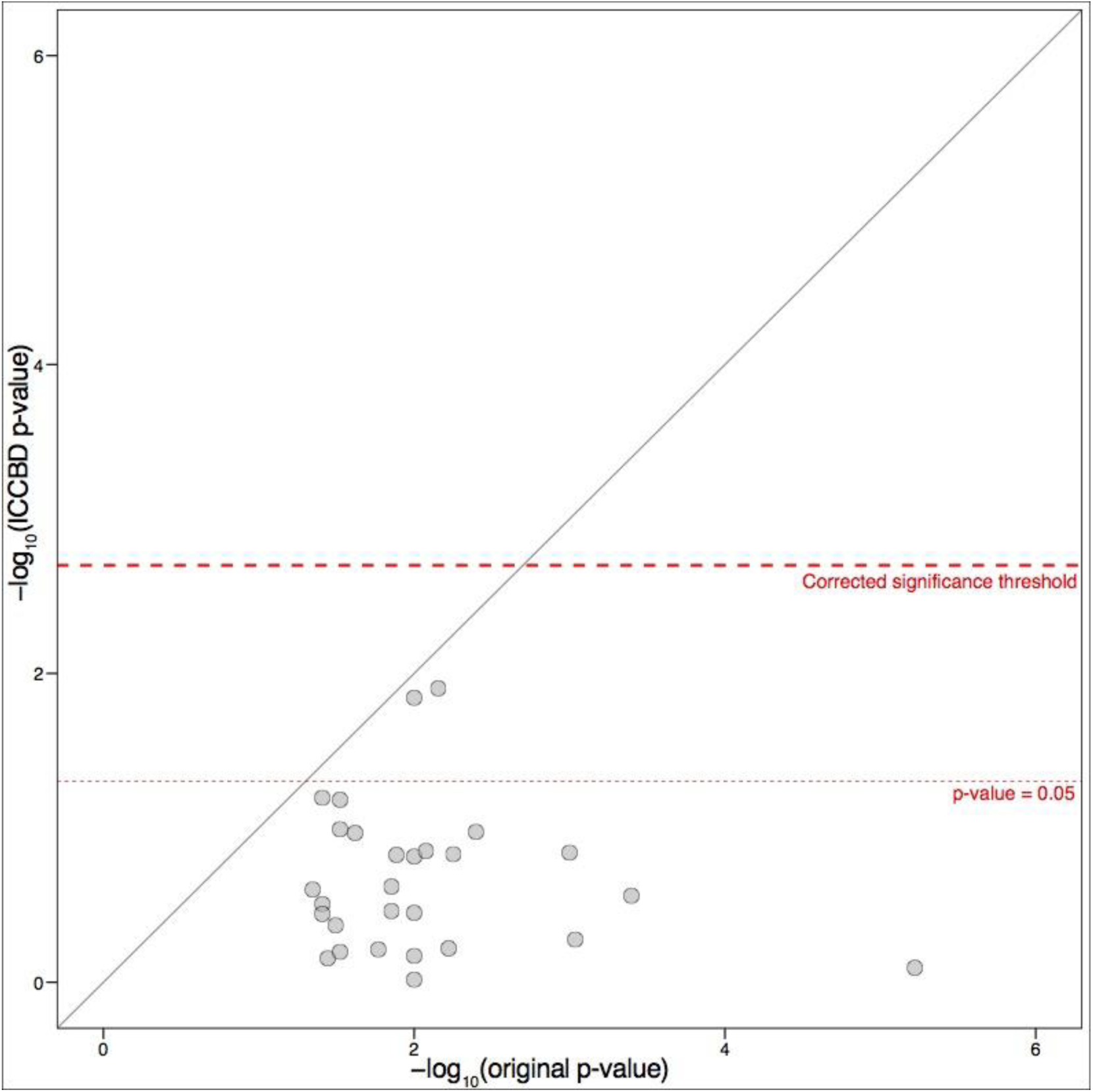
Replication of previous reports of CNV burden in BD. Curation of literature on CNV burden in BD identified 36 instances where nominal association (p-value < 0.05) was reported. We were able to test 28 of these in the ICCBD. Plotted here are p-values in previous reports (x-axis) compared to the same test performed in ICCBD cohort (y-axis). There were 4 tests for which nominal significance was observed in the ICCBD data: (1) singleton deletions greater than 100KB in cases compared to controls, (2) proportion of individuals with a singleton deletion greater than 100KB in cases compared to controls, (3) singleton deletions greater than 100KB in early onset cases compared to controls, and (4) proportion of individuals with a singleton deletion greater than 100KB in early onset cases compared to controls. None of these observations surpassed multiple test correction for the 27 tests we followed up in our data.

We next sought to assess CNVs previously implicated in psychiatric diseases for contribution to BD risk. We compiled CNVs from previous reports into 3 lists: a BD set based on *de novo* CNV events observed in BD trios^16,17,24^; a SCZ set derived from a study of over 20,000 SCZ cases and 20,000 controls^27^, and a set comprised of CNVs previously implicated in neurodevelopmental disorders^17^. The latter two sets shared a subset of regions but were treated as independent sets after accounting for the overlap (see ***Methods***). Neither the BD nor SCZ sets were enriched for deletions or duplications in cases compared to controls. A nominal enrichment that did not survive correction was noted for the neurodevelopmental set (p-value = 0.007).

BD is a heterogeneous disorder clinically, and a previous report of common variation in this cohort^12^ found evidence for genetic heterogeneity between clinical subtypes of BD. This information, combined with CNV burden being a well-established component of SCZ genetic architecture^27^, led us to hypothesize that increased CNV burden may be present in BD subtypes with high clinical similarity to SCZ. To test this hypothesis, we first sought to determine if CNV burden differed between BD subtypes (BD I n = 3,833, BD II n= 1,436, SAB n = 579) and controls (n = 6,383), as well as between BD subtypes compared to one another. Increased burden was seen in SAB compared to controls in all 3 of the primary burden classes evaluated, as well as compared to both BD I and BD II (***Table 2***). For one burden class – number of CNVs with size over 500KB and frequency < 1% – SAB had higher burden compared to controls (p-value = 0.001), BD I (p-value = 3 × 10^−4^; ***Figure 2a***) and BD II (p-value = 7 × 10^−4^). We therefore elected to focus downstream CNV analyses on this class of burden.

**Figure 2.**
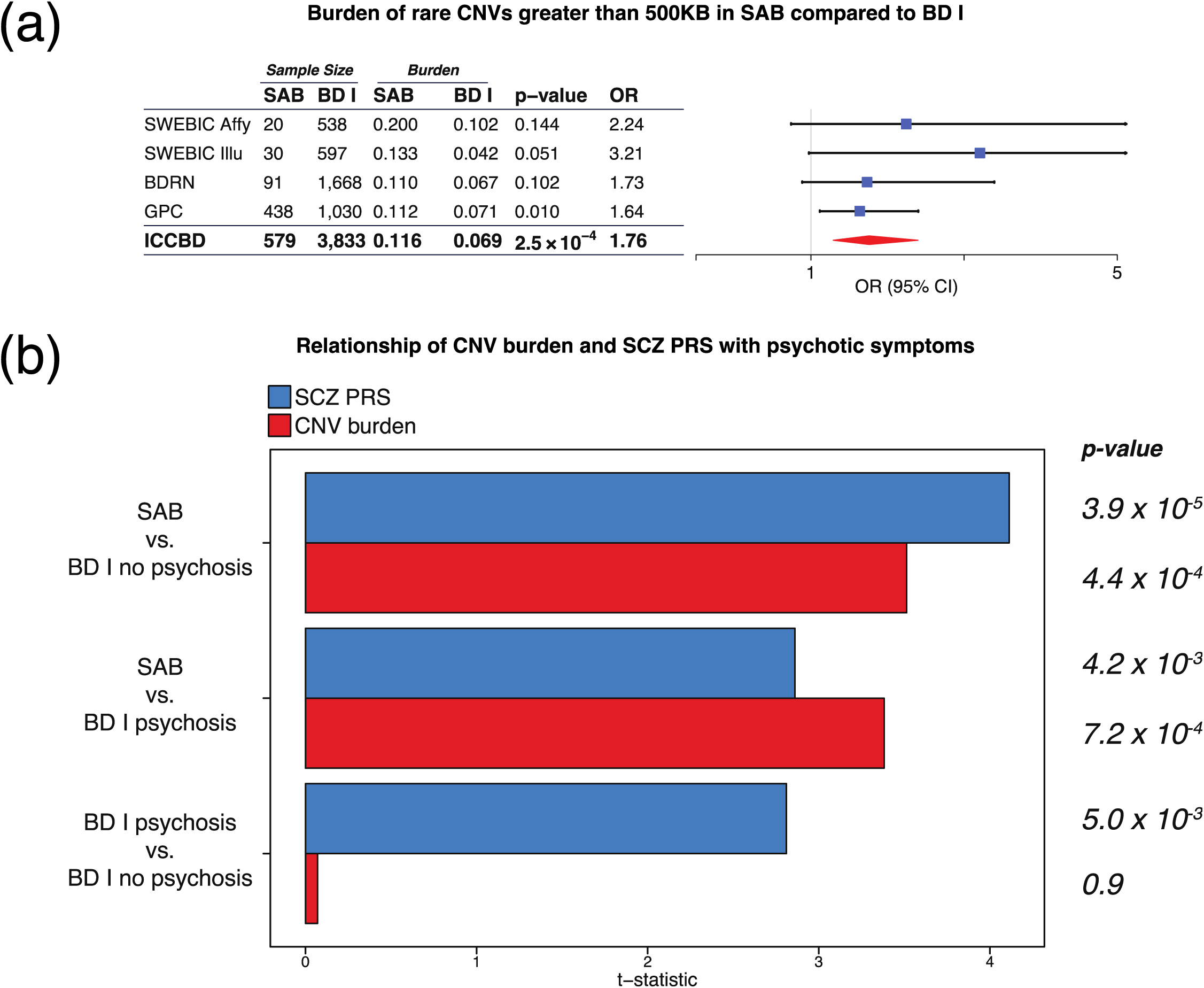
Burden of rare CNVs (frequency < 1%) greater than 500KB in SAB compared to BD I. (a) Forest plot of CNV burden partitioned by site of collection, with the full ICCBD sample at the bottom. CNV burden is calculated by combining CNV deletions and duplications. The p-values presented here for burden tests used a logistic regression model predicting SAB-BD I status by CNV burden along with covariates. The odds ratio (OR) is the exponential of the logistic regression coefficient, and OR > 1 predicts increased SAB risk. (b) Comparison of BD and SAB to one another with respect to polygenic risk scores and CNV burden. Regression analyses were performed of phenotype (stratified by history of psychosis) on polygenic scores derived from a previous GWAS for SCZ (blue) and burden of CNVs with frequency less than 1% and size greater than 500KB (red). MDS components, study site and gender were used as covariates. The t-statistic plotted on the x-axis is the ratio of the coefficient of the polygenic score or CNV burden variable and its standard error from the generalized linear model regression equation. The direction of the plotted bars indicates higher CNV burden or PRS in the phenotype listed first in the y-axis label. The p-values for whether polygenic risk scores or CNV burden differed significantly between phenotypes are shown at the far right.

### Contribution of CNV burden and SCZ PRS to psychosis in BD

SCZ is the archetypal psychotic illness in current psychiatric classification systems^5^ and increased CNV burden is a well-established component of its genetic architecture^27,28^. Psychosis is also a prominent component of BD, and the diagnostic criteria differentiating BD subtypes (e.g., BD I, SAB) from one another and from SCZ relate to the co-occurrence of psychosis with mania^5,6^. The observed CNV burden in SAB – a diagnosis that requires most of the criteria of SCZ be met – being absent in BD broadly prompted inquiry into whether CNV burden contributes to psychosis or to non-diagnostic clinical phenomena that differentiate SAB from other BD subtypes, and whether the same pattern is seen for common SCZ risk alleles. We stratified the ICCBD cases by the prominence of psychotic symptoms, correlating psychosis risk with both the CNV burden (number of CNVs with size over 500KB and frequency < 1%) and SCZ PRS^12,32^. Cases were stratified into SAB (n = 579), BD I with psychosis (n = 2,676) and BD I without psychosis (n = 850). CNV burden was increased in SAB compared to BD I with and without psychosis (SAB rate = 0.116; BD I with psychosis rate = 0.069, p-value = 7.21 × 10^−4^; BD I without psychosis rate = 0.067, p-value = 4.42 × 10^−4^), but no difference was observed between BD I with and without psychosis (p-value = 0.88; ***Figure 2b***). SCZ PRS were higher in SAB compared to BD I with psychosis (p-value = 0.004) and in BD I with psychosis compared to BD I without psychosis (p-value = 0.005; ***Figure 2b***).

## DISCUSSION

We observed no differences in the genome-wide burden of rare, large CNVs between BD cases and controls. This study had more than double the sample size used to initially identify CNV burden in SCZ, which is now well-established^27,28^. This suggests that the lack of signal in BD is not due to lack of power. We were also able follow up on most nominally significant genome-wide CNV burden results that had previously been reported with respect to BD, reproducing the original analysis with respect to phenotypes compared and the cutoffs for CNV size and frequency used in the quality control procedures. Depending on the test, our case sample size represented an increase in case sample size from the original report of 3.74-fold to 28.71-fold (***Table 3***). We did not find strong support for any of the previous observations with respect to genome-wide CNV burden in BD. Taken together, the BD case-control analyses presented here strongly suggest that rare CNV burden is not a feature of BD when treated as a single diagnostic entity.

Individuals with a diagnosis of BD comprise a clinically heterogeneous group, and the lack of CNV burden when BD is treated as a single diagnostic entity does not preclude a role of CNV burden in the pathogenesis of subsets of cases. Specifically, we hypothesized this may the case for individuals who present with psychotic symptoms in the absence of a major mood episode, given the known CNV burden in SCZ^27,28^ and the clinical overlap between SCZ and BD. Indeed, we found that cases with SAB – who by definition experience psychosis both in the presence and absence of mania – have higher rates of large, rare CNVs compared to controls and other BD subtypes.

The diagnostic criteria differentiating BD I with psychosis, SAB and SCZ from one another relate to the prominence and timing of psychotic symptoms. Through deeper analyses comparing SAB and BD I, however, we found that CNV burden was unrelated to the presence of psychosis. This was in contrast to SCZ PRS, which were higher in SAB compared to BD I with psychosis, and higher in BD I with psychosis compared to BD I without psychosis. Taken together, these results suggest that common SNPs may contribute to psychotic symptoms whereas rare CNVs may contribute to other dimensions of clinical illness within individuals with severe psychotic conditions. In this way, rare CNVs may contribute to the clinical phenomena that differentiate diagnostic categories but are not part of formal diagnostic criteria. One possibility in this regard is that CNVs may influence risk for cognitive deficits, which are more prominent in SCZ compared to BD. CNVs in disorders characterized by prominent cognitive deficits affect cognition in the general population^33^, and it will be of interest in future work to test if CNV burden is increased in BD patients who show cognitive impairments akin to those seen in SCZ^42,43^ as might be suggested by recent family-based studies^44^. Another possibility is that CNV burden increases risk for spontaneous psychosis (i.e., the psychoses of SCZ and SAB) but not psychosis secondary to severe mental stress, which it can be argued is the mechanism underlying psychosis during mania. Future studies with deeper phenotyping should aim to test these and other hypotheses.

This study has important limitations. Diagnostic misclassification of SCZ cases with SAB is possible, and while unlikely could account for the observed PRS and CNV results. For several of these analyses, sample size is a critical consideration and, as our inability to replicate most of the previous findings with respect to CNVs in BD highlighted, caution must be taken to avoid over-interpreting the results of analyses of this size.Instead, we emphasize that these findings must be followed up in larger cohorts. If replicated, they would provide support for the notion that different classes of genetic variants contribute to different classes of symptomatology in mood and psychotic syndromes. It might then be fair to inquire whether the higher CNV burden in SCZ compared to BD may be evidence not that they comprise two biologically distinct disease entities, but rather that clinicians are more likely to diagnose SCZ when a particular clinical phenomenon is present (e.g., cognitive deficits, spontaneous psychosis). These unresolved questions highlight the need for a multiscale approach to the study of mental illness, whereby integrating high-dimensional molecular and clinical data from each patient at the scale that GWAS has shown can be achieved may facilitate the development of a data-driven taxonomy.

## Acknowledgments

We are grateful for the participation of all subjects contributing to this research, and to the collection team that worked to recruit them. We acknowledge funding support from by National Institutes of Health (NIH)/National Institute of Mental Health (NIMH) grant R01MH085542 (AWC, JWS and PS), NIH/NIMH grant R01 MH106547 (JWS, PS, SAM), NIH/NIMH grant R01MH085548 (AHF, CNP, CPM, DM, DOP, DSL, ELB, HM, HN, JAK, JJR, JLS, LJF, MAE, MHR, MTP, PFB, SRM), NIH/NIMH grant K99MH101367 (PHL), the Stanley Medical Research Institute (JLM, KC, RAB, SAM), philanthropic gifts from Kent and Elizabeth Dauten and Ted and Vada Stanley (JLM, KC, RAB, SAM), the Swedish Research Council 2013-3196 (CMH), the Swedish Medical Research Council grants K2014-62X-14647-12-51 and K2010-61P-21568-01-4 (ML), the Swedish foundation for Strategic Research grant KF10-0039 (ML), the Swedish Federal Government under the LUA/ALF agreement grants ALF 20130032 and ALFGBG-142041 (ML), European Commission-Marie Curie Fellowship (AD), Wellcome Trust (IJ, KG, LAJ, LF, NC). JWS is an MGH Tepper Family Research Scholar. Work at the Icahn School of Medicine at Mount Sinai was also supported by the Institute for Genomics and Multiscale Biology and the the computational resources and staff expertise provided by Scientific Computing at the Icahn School of Medicine at Mount Sinai. The funders had no role in study design, execution, analysis or manuscript preparation.

## Supplementary Material

### Phenotyping

#### Swedish Bipolar Cohort (SWEBIC)

SWEBIC phenotype data is derived from 3 primary sources. The St. Göran Bipolar Project cohort performs phenotyping via psychiatrists (or residents in psychiatry) using a Swedish version of the Affective Disorder Evaluation that was employed in the STEP-BD study and which includes the SCID module for affective disorders. Other disorders are covered by the M.I.N.I. Neuropsychiatric interview (MINI). The BipoläR cohort is based on the Swedish quality assurance (QA) register. Phenotyping is performed by the registering physician in the QA register, and structured telephone interview is also conducted by research nurses. The Swedish study of bipolar disorder cohort (also called the “Schalling cohort”) performs phenotyping of patients by nurses and doctors manually reviewing medical charts and documenting a health interview form. Across the collection sites, age of onset was assigned based on age of first symptoms (<12 years old, 12-24, >24) and age at first diagnosis (<12 years old, 12-17, 18-24, 25-40, >40).

#### Bipolar Disorder Research Network (BDRN)

Phenotyping strategies for the BDRN cases and controls have been previously reported^1^. In brief, case participants were interviewed using the Schedules for Clinical Assessment in Neuropsychiatry (SCAN). Psychiatric and general practice case-notes, where available, were also reviewed. On the basis of these data, best-estimate lifetime diagnoses were made according to DSM-IV criteria, and key clinical variables were rated such as age at onset. In cases where there was doubt, diagnostic and clinical ratings were made by at least two members of the research team blind to each other’s rating. Age of onset was assigned based on age at first impairment (<12 years old, 12-17, 18-24, 25- 40, >40) and age at first symptoms (<12 years old, 12-17, 18-24, 25-40, >40).

#### Genomic Psychiatry Consortium (GPC)

A comprehensive description of the phenotype data collection procedures for the GPC cohort has been previously published^2^. In brief, GPC cases and controls were collected via the University of Southern California healthcare system. Using a combination of focused, direct interviews and data extraction from medical records, diagnoses and sub-phenotypes were established using the OPCRIT^3^. Age and gender-matched controls were ascertained from the University of Southern California health system and assessed using a validated screening instrument and medical records. Subphenotypes from the GPC included in this report include family history, age of onset, history of psychosis, and clinical course. Age of onset was assigned based on age of first impairment (<12 years old, 12-17, 18-24, 25-40, >40).

#### Inter-site phenotypic comparisons

As previously reported^1^, we established a Phenotype Committee including at least 1 trained clinician from each participating site in order to assess the comparability of phenotypic classification across ICCBD cohorts. Each site contributed a set of notes from cases and from individuals that did not meet criteria for bipolar disorder but did meet criteria for related mood or psychotic disorders (also known as distractors). The notes were compiled in such a way as to keep them blinded with respect to case/distractor status. Each record included the full de-identified and finalized set of diagnostic data that were used by the sites’ trained clinicians to evaluate diagnosis. Each of the Phenotype Committee members provided independent ratings of the primary variable (case vs. distractor) by reviewing the records. A quantitative analysis was conducted to determine the degree to which the committee members agree with diagnoses made by the trained clinicians. The inter-rater reliability was assessed using Fleiss’ Kappa statistic for multiple raters (p = 0.72 for the primary diagnostic variable).

### Quality Control

#### Accounting for platform differences in BDRN cases and controls

The BDRN dataset collected for the ICCBD consisted of cases only, as controls matched to these samples with respect to ancestry had previously been genotyped for a different study and were publicly available. These controls had been genotyped on the custom Illumina 1.2M chip designed for the WTCCC study, while the cases had been genotyped on either the Illumina ComboChip or the Illumina OmniExpress chip. In CNV studies, ideally the cases and controls should be genotyped on the same chip. When this is not possible, typically prior to CNV calling quality control measures can be performed to maximize the similarity between the input data with respect to the areas of the genome represented^4^. This ensures that one chip does not cover a region missing from the other chip, which despite adequate post-calling quality control could result in high-quality CNV calls in these regions. Such CNVs would then appear to be present only in one phenotype and artifactually associated with disease. For this study, it was not possible to perform these pre-calling quality control steps for BDRN because, while the cases and controls were called using the same pipeline, it was done several years apart, by which time the raw intensity files for the cases were no longer readily accessible. We observed many high-confidence CNV calls in BDRN controls that overlapped low-confidence CNV calls in BDRN cases, specifically calls failing filters for probe number and/or density (***Supplementary Figure)***. This artifact was due to these genomic regions being well covered by the genotype chip used for controls but poorly covered on the chip used for cases. In order to minimize this artifact, we implemented the following procedure. Prior to filtering based on probe number and density, we identified all BDRN CNVs that failed to meet passing criteria for at least 1 of these filters (that is, had less than 10 probes and/or a probe density less than 1 per every 20,000 bases). For each of these CNVs, we identified CNV events in other BDRN individuals that overlapped by at least 50%. We removed from analysis both (1) the CNV event failing the filter and (2) the overlapping CNVs in other BDRN individuals. In this way, regions showing poor coverage in either BDRN cases or BDRN controls were removed from both groups.

#### Plate effects

As our dataset was composed of cohorts collected from 3 countries and genotyped on 5 different chips, we sought to establish whether batch effects could play a role in CNV burden. The 15,009 samples in the final analysis were genotyped across 248 plates. We did a burden test for plate effects for rare CNVs after quality control filters had been applied. As the number of tests performed was approximately 250, at an alpha of 0.05 approximately 12 tests would be expected to be positive by chance alone. None of the plates showed a significantly higher number of CNVs per person compared to all others (data not shown).

### Supplementary Tables and Figures

**Supplementary Figure.**
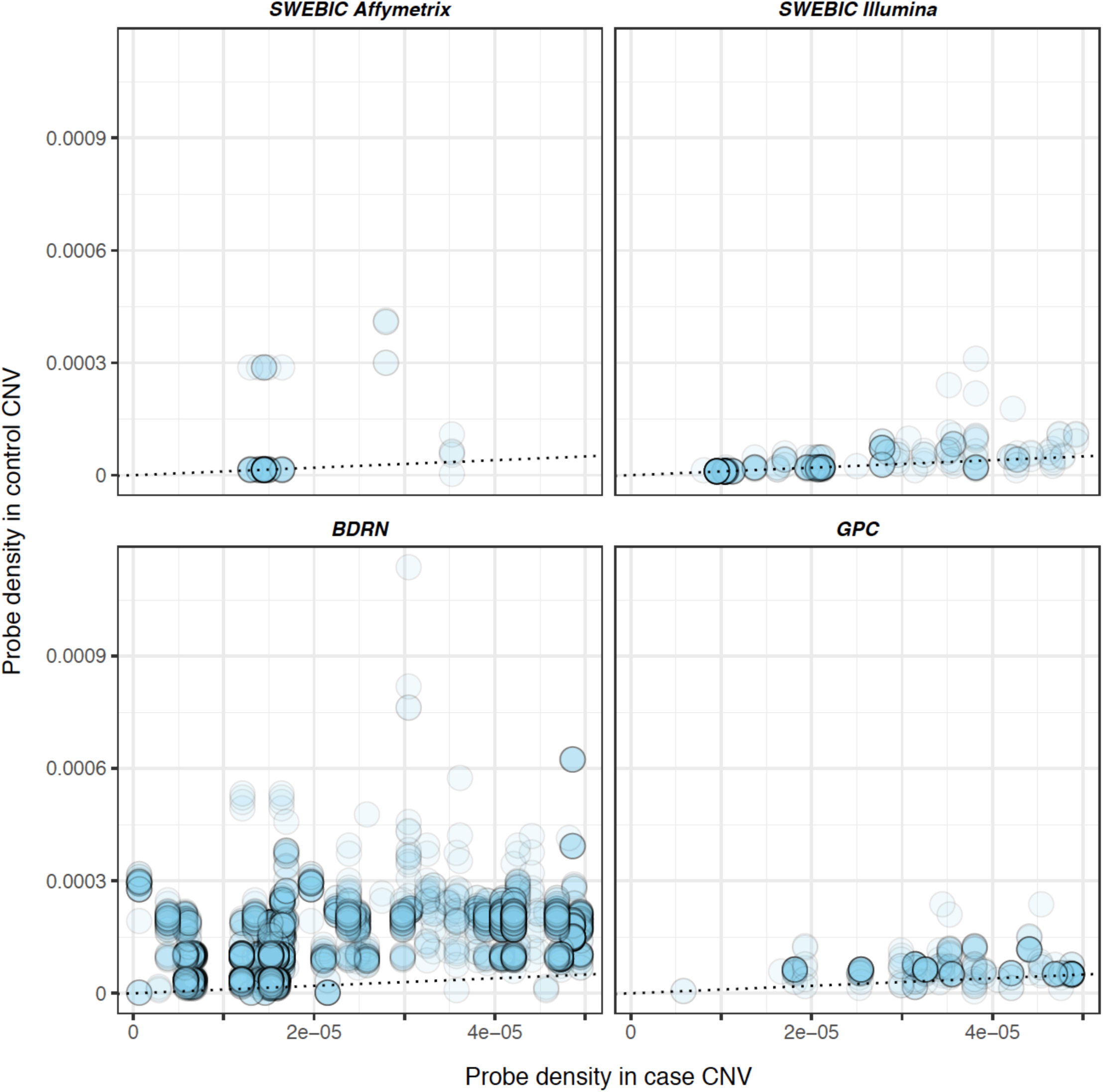
Probe density in cases compared to controls in genomic regions failing density filter in cases. For each ICCBD site, CNVs with (a) less than 1 probe per 20,000 bases and (b) overlapping any CNV event in a control from the same cohort were identified. Each point in the plot for the corresponding ICCBD site represents one such event. Plotted on the x-axis is the number of probes per base for this event in the case, and on the y-axis the number of probes per base for the overlapping CNV found in a control. In the absence of systematic differences in genomic coverage in cases and controls, we would expect points to fall on the diagonal (black dotted line), as is seen for the 3 ICCBD sites where cases and controls were genotyped on the same platform. For BDRN, the one site where cases and controls were genotyped on different platforms, we observe that controls have adequate probe coverage at sites where cases do not, which if unaccounted for would lead to artifactual enrichment of CNV rates in controls compared to cases. To account for this artifact, any CNV in a BDRN individual that overlapped a CNV from a different BDRN individual failing filter for probe number and/or density was removed.

**Supplementary Table.** Site breakdown of CNV burden in BD cases compared to controls. Results are presented for the full ICCBD cohort (6,353 cases, 8,656 controls) as well as for the ICCBD samples that remain after removing 917 cases and 1,669 controls for which genome-wide burden analyses have previously been reported. Three classes of CNV burden are presented, as designated by the description above the dotted lines. The case/control ratio is calculated by dividing the corresponding burden metric in cases by the metric in controls. P-values are two-sided, uncorrected for multiple testing, and based on 10,000 permutations testing for relative burden in cases compared to controls. CNV type, size, and frequency refer to the filters applied for the test being reported. Rows with p-values less than 0.05 in the full ICCBD case-control analysis are shown in bold. Singletons are those CNVs that occur once within the full ICCBD case-control cohort when filtered for those greater than 100KB in size. KB=kilobases

| CNV type | Size (kb) | Frequency | ICCBD |  |  |  | ICCBD without SWEBIC Affymetrix |  |  | SWEBIC Affymetrix |  | SWEBIC illumina |  | GPC |  | BDRN |  |
| --- | --- | --- | --- | --- | --- | --- | --- | --- | --- | --- | --- | --- | --- | --- | --- | --- | --- |
|  |  |  | Events | P | Case/control ratio | Control rate | P | Case/control ratio | Control rate | Case/control ratio | Control rate | Case/control ratio | Control rate | Case/control ratio | Control rate | Case/control ratio | Control rate |
| Number of CNVs per individual |  |  |  |  |  |  |  |  |  |  |  |  |  |  |  |  |  |
| All | 100 | <1% | 10515 | 0.86 | 0.99 | 0.702 | 0.62 | 0.99 | 0.658 | 1.11 | 0.89 | 1.09 | 0.61 | 0.99 | 0.68 | 0.94 | 0.66 |
|  |  | Singleton | 1849 | 0.53 | 1.03 | 0.122 | 0.62 | 1.03 | 0.137 | 1.13 | 0.14 | 1.08 | 0.10 | 1.02 | 0.13 | 1.01 | 0.12 |
|  | 500 | <1% | 1014 | 0.35 | 1.06 | 0.066 | 0.06 | 1.15 | 0.056 | 0.95 | 0.11 | 1.15 | 0.04 | 1.04 | 0.08 | 1.18 | 0.05 |
|  |  | Singleton | 270 | 0.24 | 1.16 | 0.017 | 0.27 | 1.16 | 0.019 | 1.08 | 0.02 | 1.89 | 0.01 | 1.10 | 0.02 | 1.18 | 0.02 |
| Deletions | 100 | <1% | 3970 | 0.78 | 1.01 | 0.264 | 0.85 | 1.01 | 0.250 | 1.09 | 0.32 | 1.14 | 0.24 | 0.91 | 0.28 | 0.98 | 0.24 |
|  |  | Singleton | 918 | 0.02 | 1.17 | 0.057 | 0.17 | 1.12 | 0.062 | 1.42 | 0.06 | 1.25 | 0.05 | 1.10 | 0.07 | 1.06 | 0.05 |
|  | 500 | <1% | 231 | 0.11 | 1.24 | 0.014 | 0.17 | 1.27 | 0.012 | 1.35 | 0.02 | 1.15 | 0.02 | 1.41 | 0.01 | 1.11 | 0.01 |
|  |  | Singleton | 107 | 0.06 | 1.44 | 0.006 | 0.23 | 1.35 | 0.006 | 2.24 | 0.01 | 1.05 | 0.01 | 1.18 | 0.01 | 1.27 | 0.00 |
| Duplications | 100 | <1% | 6545 | 0.72 | 0.99 | 0.439 | 0.44 | 0.98 | 0.408 | 1.12 | 0.57 | 1.07 | 0.37 | 1.06 | 0.40 | 0.92 | 0.42 |
|  |  | Singleton | 1442 | 0.57 | 0.97 | 0.098 | 0.64 | 0.97 | 0.105 | 1.15 | 0.12 | 0.94 | 0.08 | 1.01 | 0.10 | 0.92 | 0.10 |
|  | 500 | <1% | 783 | 0.85 | 1.01 | 0.052 | 0.17 | 1.12 | 0.044 | 0.84 | 0.08 | 1.15 | 0.03 | 0.97 | 0.07 | 1.19 | 0.04 |
|  |  | Singleton | 226 | 1.00 | 1.01 | 0.015 | 0.61 | 1.08 | 0.015 | 0.93 | 0.02 | 3.08 | 0.00 | 1.04 | 0.02 | 1.02 | 0.01 |
| Number of genes within CNVs per individual |  |  |  |  |  |  |  |  |  |  |  |  |  |  |  |  |  |
| All | 100 | <1% | 10515 | 0.66 | 0.97 | 1.517 | 0.19 | 0.94 | 1.330 | 1.21 | 2.30 | 1.19 | 1.09 | 1.11 | 1.24 | 0.81 | 1.41 |
|  |  | Singleton | 1849 | 0.37 | 1.10 | 0.303 | 1.00 | 1.00 | 0.330 | 1.74 | 0.39 | 1.40 | 0.20 | 1.05 | 0.27 | 0.87 | 0.31 |
|  | 500 | <1% | 1014 | 0.09 | 1.22 | 0.329 | 0.17 | 1.18 | 0.271 | 1.54 | 0.58 | 1.44 | 0.12 | 1.12 | 0.39 | 1.19 | 0.27 |
|  |  | Singleton | 270 | 0.05 | 1.51 | 0.082 | 0.14 | 1.32 | 0.085 | 3.13 | 0.11 | 3.04 | 0.03 | 0.97 | 0.10 | 1.09 | 0.08 |
| Deletions | 100 | <1% | 3970 | 0.06 | 1.15 | 0.318 | 0.31 | 1.11 | 0.262 | 1.46 | 0.55 | 1.25 | 0.32 | 1.12 | 0.28 | 0.91 | 0.24 |
|  |  | Singleton | 918 | 0.01 | 1.49 | 0.088 | 0.17 | 1.24 | 0.086 | 2.50 | 0.15 | 1.24 | 0.08 | 1.20 | 0.09 | 1.17 | 0.07 |
|  | 500 | <1% | 231 | 0.01 | 1.88 | 0.048 | 0.07 | 1.73 | 0.035 | 2.58 | 0.10 | 1.44 | 0.04 | 2.80 | 0.03 | 1.20 | 0.03 |
|  |  | Singleton | 107 | 0.02 | 2.43 | 0.021 | 0.07 | 2.35 | 0.012 | 3.88 | 0.06 | 0.96 | 0.02 | 2.23 | 0.02 | 1.57 | 0.01 |
| Duplications | 100 | <1% | 6545 | 0.28 | 0.92 | 1.199 | 0.09 | 0.90 | 1.068 | 1.12 | 1.75 | 1.17 | 0.77 | 1.11 | 0.96 | 0.79 | 1.17 |
|  |  | Singleton | 1442 | 0.87 | 0.98 | 0.277 | 0.42 | 0.92 | 0.286 | 1.57 | 0.36 | 1.30 | 0.16 | 0.96 | 0.23 | 0.80 | 0.29 |
|  | 500 | <1% | 783 | 0.45 | 1.11 | 0.281 | 0.49 | 1.10 | 0.236 | 1.31 | 0.47 | 1.44 | 0.08 | 0.96 | 0.36 | 1.19 | 0.24 |
|  |  | Singleton | 226 | 0.25 | 1.33 | 0.080 | 0.32 | 1.23 | 0.077 | 2.37 | 0.11 | 4.99 | 0.01 | 0.75 | 0.09 | 1.12 | 0.08 |
| Total distance covered by CNVs per individual |  |  |  |  |  |  |  |  |  |  |  |  |  |  |  |  |  |
| All | 100 | <1% | 10515 | 0.10 | 1.04 | 371.9 | 0.04 | 1.06 | 345.4 | 1.06 | 462.70 | 1.03 | 329.5 | 1.04 | 391.30 | 1.05 | 336.00 |
|  |  | Singleton | 1849 | 0.07 | 1.14 | 326.9 | 0.07 | 1.11 | 320.0 | 1.19 | 369.50 | 1.03 | 315.5 | 1.08 | 350.40 | 1.17 | 302.20 |
|  | 500 | <1% | 1014 | 0.60 | 1.03 | 911.0 | 0.30 | 1.04 | 838.0 | 1.08 | 1081.00 | 0.91 | 821.4 | 1.01 | 920.00 | 1.09 | 808.40 |
|  |  | Singleton | 270 | 0.34 | 1.13 | 954.6 | 0.67 | 1.04 | 913.1 | 1.43 | 1106.00 | 0.59 | 1501 | 0.93 | 1021.00 | 1.26 | 795.40 |
| Deletions | 100 | <1% | 3970 | 0.06 | 1.07 | 256.2 | 0.21 | 1.06 | 245.7 | 1.18 | 291.20 | 0.97 | 265.6 | 1.13 | 257.30 | 1.02 | 237.20 |
|  |  | Singleton | 918 | 0.21 | 1.12 | 288.8 | 0.55 | 1.06 | 271.6 | 1.39 | 321.40 | 0.67 | 392.2 | 1.20 | 273.30 | 1.09 | 259.10 |
|  | 500 | <1% | 231 | 0.84 | 1.02 | 937.5 | 0.96 | 1.01 | 891.3 | 1.07 | 1043.00 | 0.79 | 920 | 1.16 | 940.80 | 1.01 | 864.50 |
|  |  | Singleton | 107 | 0.90 | 1.02 | 1052.0 | 0.74 | 0.95 | 1025.0 | 1.14 | 1132.00 | 0.54 | 1341 | 1.16 | 951.10 | 1.09 | 940.70 |
| Duplications | 100 | <1% | 6545 | 0.35 | 1.03 | 367.7 | 0.05 | 1.06 | 342.7 | 0.98 | 447.50 | 1.05 | 306.3 | 1.03 | 396.80 | 1.07 | 336.00 |
|  |  | Singleton | 1442 | 0.15 | 1.13 | 344.0 | 0.04 | 1.13 | 326.4 | 1.10 | 429.10 | 1.35 | 255.9 | 0.99 | 377.40 | 1.18 | 314.20 |
|  | 500 | <1% | 783 | 0.59 | 1.03 | 893.3 | 0.25 | 1.05 | 818.1 | 1.12 | 1067.00 | 0.98 | 758.9 | 0.97 | 904.90 | 1.12 | 790.20 |
|  |  | Singleton | 226 | 0.32 | 1.16 | 946.4 | 0.25 | 1.11 | 848.3 | 1.43 | 1181.00 | 0.78 | 1230 | 0.84 | 1037.00 | 1.29 | 774.40 |

## References

1. Dunayevich, E. & Keck, P. E. Prevalence and description of psychotic features in bipolar mania. Curr. Psychiatry Rep. 2, 286–290 (2000).

2. Burdick, K. E., Ketter, T. A., Goldberg, J. F. & Calabrese, J. R. Assessing cognitive function in bipolar disorder: challenges and recommendations for clinical trial design. J. Clin. Psychiatry 76, e342–50 (2015).

3. Merikangas, K. R. et al. Lifetime and 12-Month Prevalence of Bipolar Spectrum Disorder in the National Comorbidity Survey Replication. Arch. Gen. Psychiatry 64, 543–552 (2007).

4. Perlis, R. H. et al. Predictors of Recurrence in Bipolar Disorder: Primary Outcomes From the Systematic Treatment Enhancement Program for Bipolar Disorder (STEP-BD). Am. J. Psychiatry 163, 217–224 (2006).

5. American Psychiatric Association. Diagnostic and statistical manual of mental disorders, Fifth Edition. (American Psychiatric Pub, 2013). doi:10.1176/appi.books.9780890425596.744053

6. World Health Organization. The ICD-10 Classification of Mental and Behavioural Disorders. Int. Classif. 10, 1–267 (1992).

7. American Psychiatric Association. Diagnostic and Statistical Manual of Mental Disorders, Third Edition. (1980).

8. American Psychiatric Association. Diagnostic and statistical manual of mental disorders, Fourth Edition, Text Revision. (Fourth edition. Washington, DC?: American Psychiatric Association, [1994] (c)1994, 2000).

9. Stahl, E. et al. Genomewide association study identifies 30 loci associated with bipolar disorder. bioRxiv (2017). doi:10.1101/173062

10. Bulik-Sullivan, B. et al. An atlas of genetic correlations across human diseases and traits. Nat Genet 47, 1236–1241 (2015).

11. Lee, S. H. et al. Genetic relationship between five psychiatric disorders estimated from genome-wide SNPs. Nat Genet 45, 984–994 (2013).

12. Charney, A. W. et al. Evidence for genetic heterogeneity between clinical subtypes of bipolar disorder. Transl. Psychiatry (in press), (2016).

13. Green, E. K. et al. Copy number variation in bipolar disorder. Mol. Psychiatry 21, 89–93 (2016).

14. Yang, S. et al. Genomic landscape of a three-generation pedigree segregating affective disorder. PLoS One 4, 1–10 (2009).

15. Zhang, D. et al. Singleton deletions throughout the genome increase risk of bipolar disorder. Mol Psychiatry 14, 376–380 (2009).

16. Malhotra, D. et al. High frequencies of de novo CNVs in bipolar disorder and schizophrenia. Neuron 72, 951–963 (2011).

17. Georgieva, L. et al. De novo CNVs in bipolar affective disorder and schizophrenia. Hum. Mol. Genet. 23, 6677–6683 (2014).

18. Chen, Y.-H. et al. Identifying Potential Regions of Copy Number Variation for Bipolar Disorder. Microarrays 3, 52–71 (2014).

19. Chen, J. et al. A pilot study on commonality and specificity of copy number variants in schizophrenia and bipolar disorder. Transl. Psychiatry 6, e824 (2016).

20. Bergen, S. E. et al. Genome-wide association study in a Swedish population yields support for greater CNV and MHC involvement in schizophrenia compared with bipolar disorder. Mol. Psychiatry 17, 880 (2012).

21. McQuillin, A. et al. Analysis of genetic deletions and duplications in the University College London Bipolar Disorder case control sample. Eur. J. Hum. Genet. 19, 588–592 (2011).

22. Grozeva, D. et al. Rare copy number variants: a point of rarity in genetic risk for bipolar disorder and schizophrenia. Arch Gen Psychiatry 67, 318–327 (2010).

23. Grozeva, D. et al. Reduced burden of very large and rare CNVs in bipolar affective disorder. Bipolar Disord. 15, 893–898 (2013).

24. Noor, A. et al. Copy number variant study of bipolar disorder in Canadian and UK populations implicates synaptic genes. Am. J. Med. Genet. Part B Neuropsychiatr. Genet. 165, 303–313 (2014).

25. Malhotra, D. & Sebat, J. CNVs: harbingers of a rare variant revolution in psychiatric genetics. Cell 148, 1223–1241 (2012).

26. Priebe, L. et al. Genome-wide survey implicates the influence of copy number variants (CNVs) in the development of early-onset bipolar disorder. Mol. Psychiatry 17, 421–432 (2012).

27. Marshall, C. R. et al. Contribution of copy number variants to schizophrenia from a genome-wide study of 41,321 subjects. Nat. Genet. 49, 27–35 (2016).

28. International Schizophrenia Consortium. Rare chromosomal deletions and duplications increase risk of schizophrenia. Nature 455, 237–241 (2008).

29. Green, E. K. et al. Replication of bipolar disorder susceptibility alleles and identification of two novel genome-wide significant associations in a new bipolar disorder case-control sample. Mol Psychiatry 18, 1302–1307 (2013).

30. McCarthy, S. E. et al. Microduplications of 16p11.2 are associated with schizophrenia. Nat. Genet. 41, 1223–7 (2009).

31. Ruderfer, D. M. et al. Polygenic dissection of diagnosis and clinical dimensions of bipolar disorder and schizophrenia. Mol Psychiatry 19, 1017–1024 (2014).

32. Bipolar Disorder and Schizophrenia Working Group of the Psychiatric Genomics Consortium. Genomic Dissection of Bipolar Disorder and Schizophrenia, Including 28 Subphenotypes. Cell 173, 1705–1715.e16 (2018).

33. Stefansson, H. et al. CNVs conferring risk of autism or schizophrenia affect cognition in controls. Nature 361–366 (2013). doi:10.1038/nature12818

34. Kraepelin, E. Manic-depressive insanity and paranoia. (E. & S. Livingstone, 1921).

35. Kraepelin, E. & Barclay, R. (translator). Dementia Praecox and the Paraphrenia. (E & S Livingstone, 1919).

36. Kendall, K. M. et al. Cognitive Performance Among Carriers of Pathogenic Copy Number Variants: Analysis of 152,000 UK Biobank Subjects. Biol. Psychiatry 103–110 (2016). doi:10.1016/j.biopsych.2016.08.014

37. Pato, M. T. et al. The genomic psychiatry cohort: partners in discovery. Am. J. Med. Genet. B. Neuropsychiatr. Genet. 162, 306–12 (2013).

38. Korn, J. M. et al. Integrated genotype calling and association analysis of SNPs, common copy number polymorphisms and rare CNVs. Nat. Genet. 40, 1253 (2008).

39. Purcell, S. et al. PLINK: a tool set for whole-genome association and population-based linkage analyses. Am. J. Hum. Genet. 81, 559 (2007).

40. International Schizophrenia Consortium. Common polygenic variation contributes to risk of schizophrenia and bipolar disorder. Nature 460, 748 (2009).

41. Schizophrenia Psychiatric Genome-Wide Association Study Consortium. Genome-wide association study identifies five new schizophrenia loci. Nat Genet 43, 969–976 (2011).

42. Burdick, K. E. et al. Empirical evidence for discrete neurocognitive subgroups in bipolar disorder: clinical implications. Psychol. Med. 44, 3083–3096 (2014).

43. Van Rheenen, T. E. et al. Characterizing cognitive heterogeneity on the schizophrenia–bipolar disorder spectrum. Psychol. Med. 47, 1848–1864 (2017).

44. Russo, M. et al. Neurocognitive subtypes in patients with bipolar disorder and their unaffected siblings. Psychol. Med. 47, 2892–2905 (2017).

## References

1 Charney AW, Ruderfer DM, Stahl EA, Moran JL, Chambert K, Belliveau RA et al. Evidence for genetic heterogeneity between clinical subtypes of bipolar disorder. Transl Psychiatry 2016; 7: e993–e993.

2 Pato MT, Sobell JL, Medeiros H, Abbott C, Sklar BM, Buckley PF et al. The genomic psychiatry cohort: partners in discovery. Am J Med Genet B Neuropsychiatr Genet 2013; 162: 306–12.

3 McGuffin P, Farmer A, Harvey I. A polydiagnostic application of operational criteria in studies of psychotic illness: development and reliability of the OPCRIT system. Arch Gen Psychiatry 1991; 48: 764.

4 Marshall CR, Howrigan DP, Merico D, Thiruvahindrapuram B, Wu W, Greer DS et al. Contribution of copy number variants to schizophrenia from a genome-wide study of 41,321 subjects. Nat Genet 2016; 49: 27–35.

